# Assessing large multimodal models for one-shot learning and interpretability in biomedical image classification

**DOI:** 10.1101/2023.12.31.573796

**Authors:** Wenpin Hou, Qi Liu, Huifang Ma, Yilong Qu, Zhicheng Ji

## Abstract

Image classification plays a pivotal role in analyzing biomedical images, serving as a cornerstone for both biological research and clinical diagnostics. We demonstrate that large multimodal models (LMMs), like GPT-4, excel in one-shot learning, generalization, interpretability, and text-driven image classification across diverse biomedical tasks. These tasks include the classification of tissues, cell types, cellular states, and disease status. LMMs stand out from traditional single-modal classification approaches, which often require large training datasets and offer limited interpretability.

## Introductions

Large multimodal models (LMMs)^1–5^ are an advanced extension of large language models (LLMs), incorporating multi-sensory capabilities to process and integrate diverse data modalities such as text, images, and audio. The ability to incorporate natural language makes LMMs distinct from single-modal methods that are built and trained solely on images. Although LMMs may not fully understand concepts in natural language, they can link these concepts with features in images and use them as tools to efficiently capture and convey information. In contrast, single-modal methods require representing features in images with a large number of parameters, and the training process often involves updating millions of parameters in a complex neural network.

We hypothesize that LMMs outperform conventional single-modal deep-learning methods in biomedical imaging classification, offering advances in *post hoc* interpretability and higher accuracy. Image classification is a classic application of deep learning methods, but the lack of interpretability has hindered the widespread application of these “black boxes” in the biomedical field. In contrast, LMMs, inheriting language capabilities from LLMs, can provide reasoning steps for image classification in plain language. This *post hoc* interpretability is crucial for building trust, ensuring transparency, and identifying errors in clinical decision-making. To test this *post hoc* interpretability, we design one-shot learning tasks in this study to evaluate and compare the performance of LMMs and single-modal models. “One-shot,” rather than “few-shot,” was chosen to highlight the application for rare diseases, where collecting, annotating, and curating training samples can be time-consuming, labor-intensive, or even inaccessible. Compared to single-modal methods, which often require vast amounts of training images, pretrained LMMs have demonstrated superior performance in one-shot learning^3^, where only one example is used in the training process. This approach eliminates the need for large training datasets, significantly reducing the complexity of applying these methods. Additionally, biomedical images generated by different labs, under varying conditions, or with differing experimental procedures, are often not directly comparable, leading to discrepancies between training and testing datasets. LMMs could be better suited to handle such discrepancies, as concepts expressed in natural language are more abstract and can generalize across images to capture common features of interest. In contrast, single-modal methods are more vulnerable to the challenges posed by dataset shifts. Finally, LMMs support in-context learning, where users can directly provide training examples via prompt messages without needing to retrain or fine-tune the model. Since several commercial LMMs offer user-friendly interfaces that require no programming skills, they have the potential to significantly lower the barrier to performing biomedical image classification tasks, improving health equity and benefiting hard-to-reach populations and resource-limited areas. By contrast, implementing single-modal methods often requires substantial programming skills and deep learning expertise.

Previous studies have explored the application of LMMs in biomedical image analysis to a limited extent^3,6,7^, but they fail to address key aspects of biomedical image classification. These studies either neglect comparisons with single-modal methods^3,6^ or include only a small subset of them^7^, leaving the advantages of LMMs over single-modal approaches largely unclear. Additionally, they primarily rely on simple prompt strategies without exploring alternative approaches that could be applied in diverse scenarios. The natural language outputs generated by LMMs are evaluated qualitatively and descriptively, but a systematic assessment of their accuracy in describing images is missing, which is crucial for understanding the interpretability of LMMs. Moreover, these studies largely focus on zero-shot performance, where no training data is provided. However, recent LMMs often avoid providing definitive answers in zero-shot settings, likely due to legal concerns related to misclassification, making zero-shot performance less relevant in practical applications.

In this study, we addressed these limitations in previous studies by systematically benchmarking the performance of several popular LMMs, including GPT-4o, Claude 3.5, Gemini 1.5, and Llama 3.2, across eight types of biomedical image classification tasks. Our evaluation focused on the one-shot learning ability and interpretability of LMMs, which we believe are the aspects that most clearly distinguish LMMs from single-modal methods. To test the generalizability of LMMs, we introduced discrepancies between the training and testing data by applying various transformations to the training images. For comparison, we repeated all analyses using ten conventional single-modal image classification methods, four single-modal image classification methods focused on few-shot learning, and three image captioning methods. Our findings show that LMMs substantially outperform other methods in one-shot learning, interpretability, and generalizability, highlighting the unique advantages of LMMs in biomedical image classification and offering new insights not reported in previous benchmark studies^6,8,9^.

## Results

### Prompt strategies for image classification

We developed three prompt strategies for an LMM to perform image classification. In the first strategy (Figure 1a), a training image array with two rows and two columns is created for one-shot binary classification tasks. The two rows represent two image categories, and the first column labels these as “row 1” and “row 2.” The second column contains two training images, one for each category. The training image array can easily be created using PowerPoint, Keynote, or similar software. A dictionary of tissues, cell types, or disease conditions corresponding to the two rows is stored on the user’s local system and is hidden from the LMM. When presented with a test image, the LMM is asked to identify which row in the training image array most closely resembles the test image. The user can then refer to the locally stored dictionary to determine the actual category.

**Figure 1.**
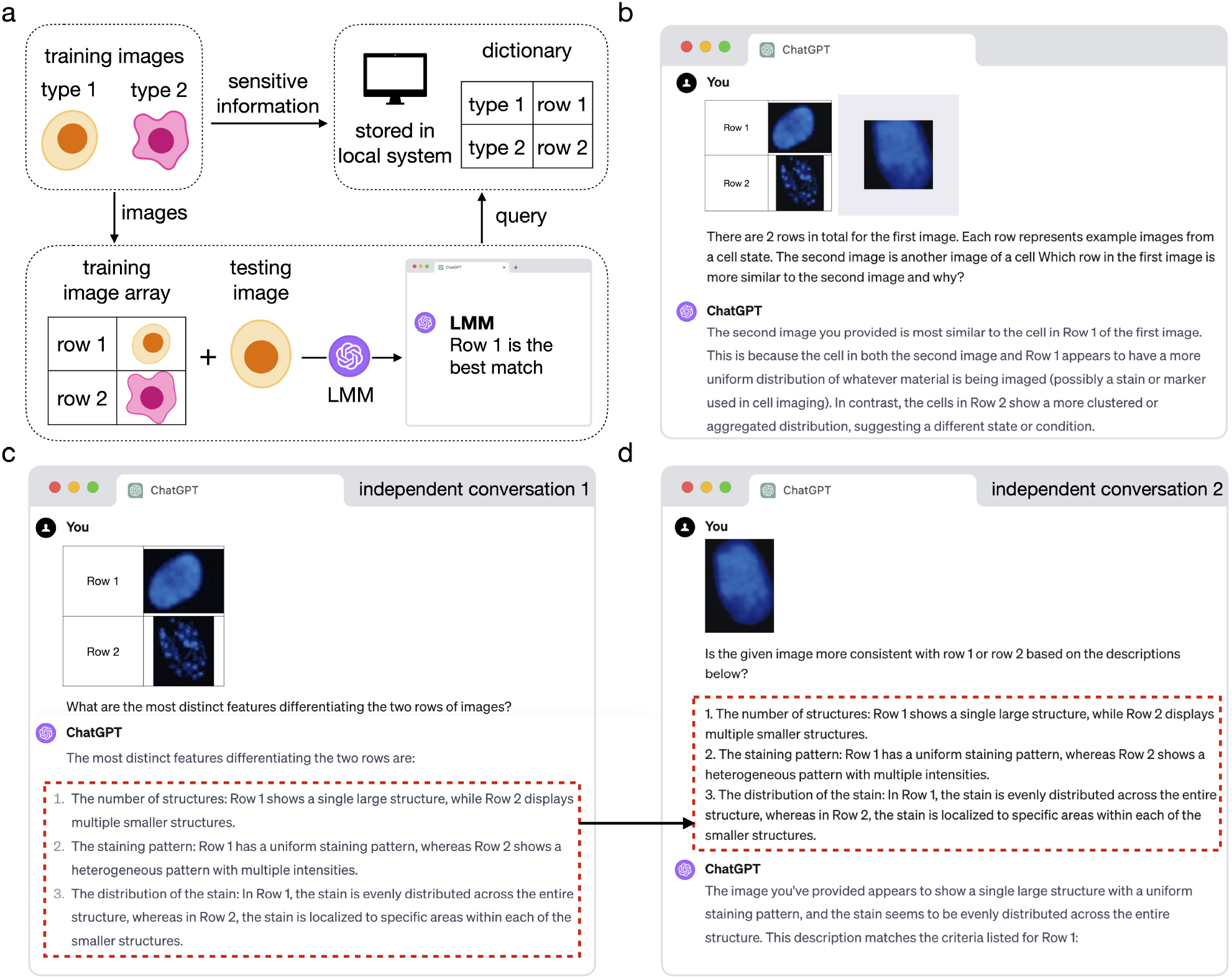
Demonstration of LMM image classification. **a**, A schematic of the prompt strategy for LMM image-based classification. **b**, An example conversation with GPT-4 performing image-based classification. **c, d**, An example process with GPT-4 for text-based classification. The process consists of two independent conversations: one to obtain a text description of the training image’s distinct features (**c**) and another to query GPT-4 for classifying a test image (**d**).

This strategy offers several advantages. First, additional rows and columns can be added to represent more image categories or include more training images per category, thus bypassing the maximum input file limits imposed by many LMMs. Second, compared to providing separate image files for each category, the reference image array simplifies communication for both the user and the LMM. Third, sensitive information, such as patient data, can be stored in a separate dictionary for enhanced confidentiality, which is especially important in clinical settings. Finally, this design minimizes the chance that the LMM will rely on prior knowledge of image category names (e.g., cell type names), ensuring a fair comparison with single-modal methods. Figure 1b illustrates a real example where GPT-4 was queried to classify images of normal and senescent cells using this prompt strategy.

In addition, we designed a second text-based image classification strategy that uses the LMMs’ natural language output from a previous conversation as training information in a completely new conversation (Figure 1c-d). First, the LMM was asked to summarize the main differences between the two image categories in the training image array (Figure 1c). Then, in a new conversation, the LMM was queried to identify the category of a test image using only the summarized text, without access to the original training images (Figure 1d). We also designed a third strategy in which the text description of each training and testing image was converted into a numeric vector of text embeddings, and classification was performed by comparing these embeddings using cosine similarity (Methods).

### LMMs outperform single-modal methods in one-shot learning

We systematically evaluated the performance of four LMMs (GPT-4o, Claude 3.5 Sonnet, Gemini 1.5 Pro, and Llama 3.2 11B vision) for in-context one-shot learning, with one training image provided for each image category. Additionally, we included ten conventional single-modal methods and four few-shot single-modal methods, which were benchmarked on the same sets of training and testing images.

We designed four biology image classification tasks and four medical image classification tasks (Figure 2). The biology image tasks include classifying artery and tibial nerve tissues using hematoxylin and eosin stain (H&E) images, classifying artery, tibial nerve, and adipose tissues using H&E images, classifying microglial (BV-2 cell line) and neuroblastoma (SH-SY5Y cell line) cells using light microscopy images, and classifying normal and senescent cells using DAPI fluorescent stain images. The medical image tasks include classifying normal retinas and retinas with diabetic macular edema (DME) using optical coherence tomography (OCT) images, classifying normal brains and brains with tumors using magnetic resonance imaging (MRI), classifying normal lungs and lungs with tumors using chest computed tomography (CT) scans, and classifying H&E histology images with high or low density of glioblastoma (GBM) cancer cells.

**Figure 2.**
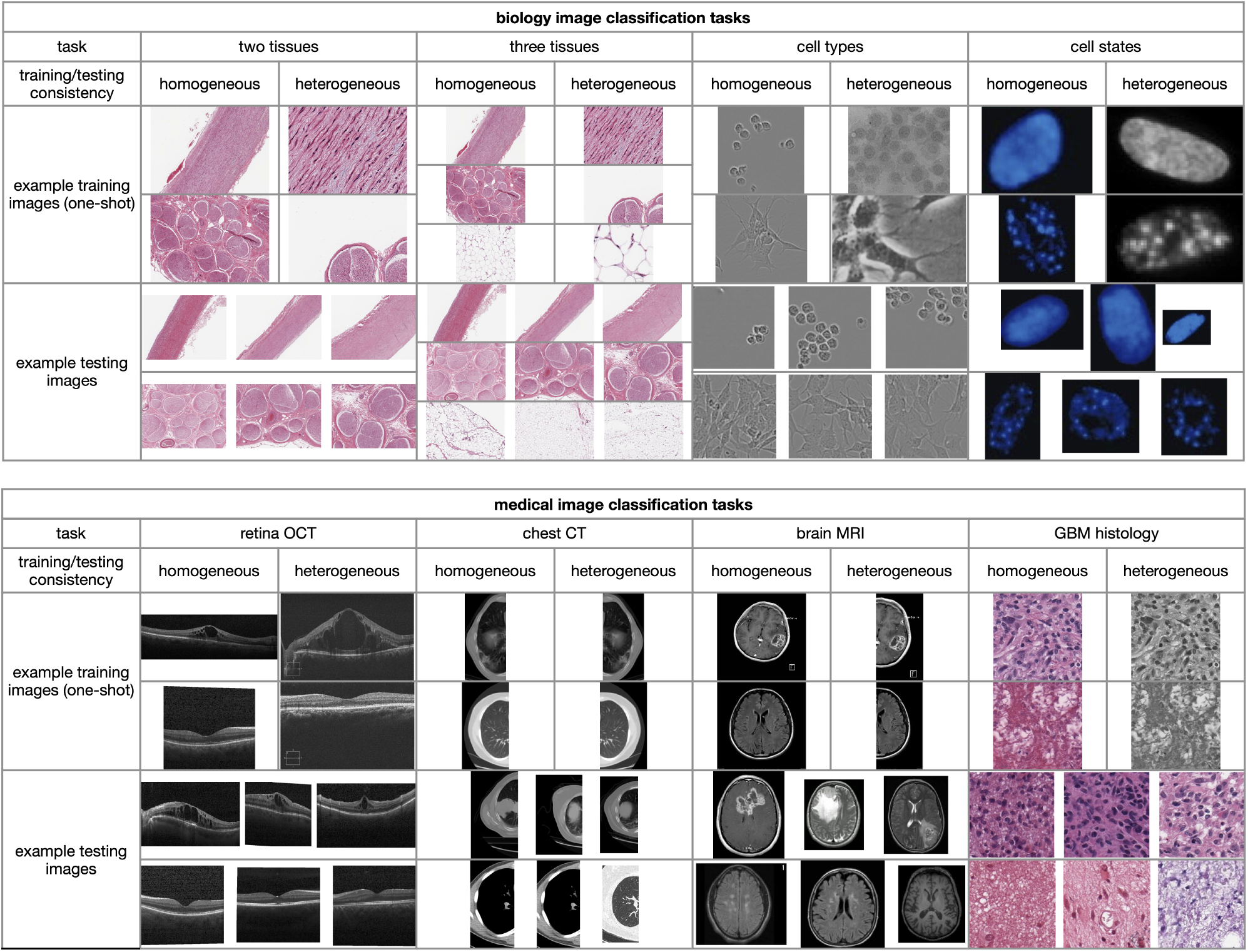
Example training and testing images for four biology and four medical image classification tasks. The image categories in each dataset, from top to bottom, are as follows: artery and tibial nerve for two tissues; artery, tibial nerve, and adipose for three tissues; BV-2 and SH-SY5Y for cell types; normal and senescent cells for cell states; DME and normal for retina OCT; cancer and normal for chest CT; cancer and normal for brain MRI; and high and low GBM cancer cell density for GBM histology.

Figure 3 shows the accuracy, defined as the percentage of test images correctly categorized (Methods), of each method across the classification tasks. The accuracy and responses of the LMMs are detailed in Supplementary Table 1. GPT-4o (labeled as “GPT-4o image” in Figure 3) demonstrates the best performance, achieving nearly 100% accuracy in all scenarios. The other two LMMs, Claude 3.5 and Gemini 1.5, also show strong performance in most scenarios but do not match the performance of GPT-4o. The ten conventional single-modal methods and four few-shot single-modal methods perform worse than the three LMMs. For instance, SwinTransformer, the best-performing conventional single-modal method, performs significantly worse than GPT-4o in the four medical image tasks, despite achieving near-perfect performance in the biology image tasks. Similarly, Self+RestoreNet, the best-performing few-shot single-modal method, also struggled to perform well in two medical imaging tasks. A potential explanation is that the differences between the two image categories in the medical tasks are smaller and more subtle compared to those in the biology tasks. As a result, single-modal methods may require more training data to effectively capture such differences. In contrast, GPT-4o is able to capture these differences using only one training image per category. Another open-source LMM, Llama 3.2 11B Vision, has the worst performance among all these methods, with results that are close to random guessing in almost all tasks. A potential reason for this is that Llama 3.2 11B Vision has a substantially smaller number of model parameters compared to GPT-4o, Claude 3.5 Sonnet, and Gemini 1.5 Pro, thereby lacking the capacity to accurately differentiate between different categories of images.

**Figure 3.**
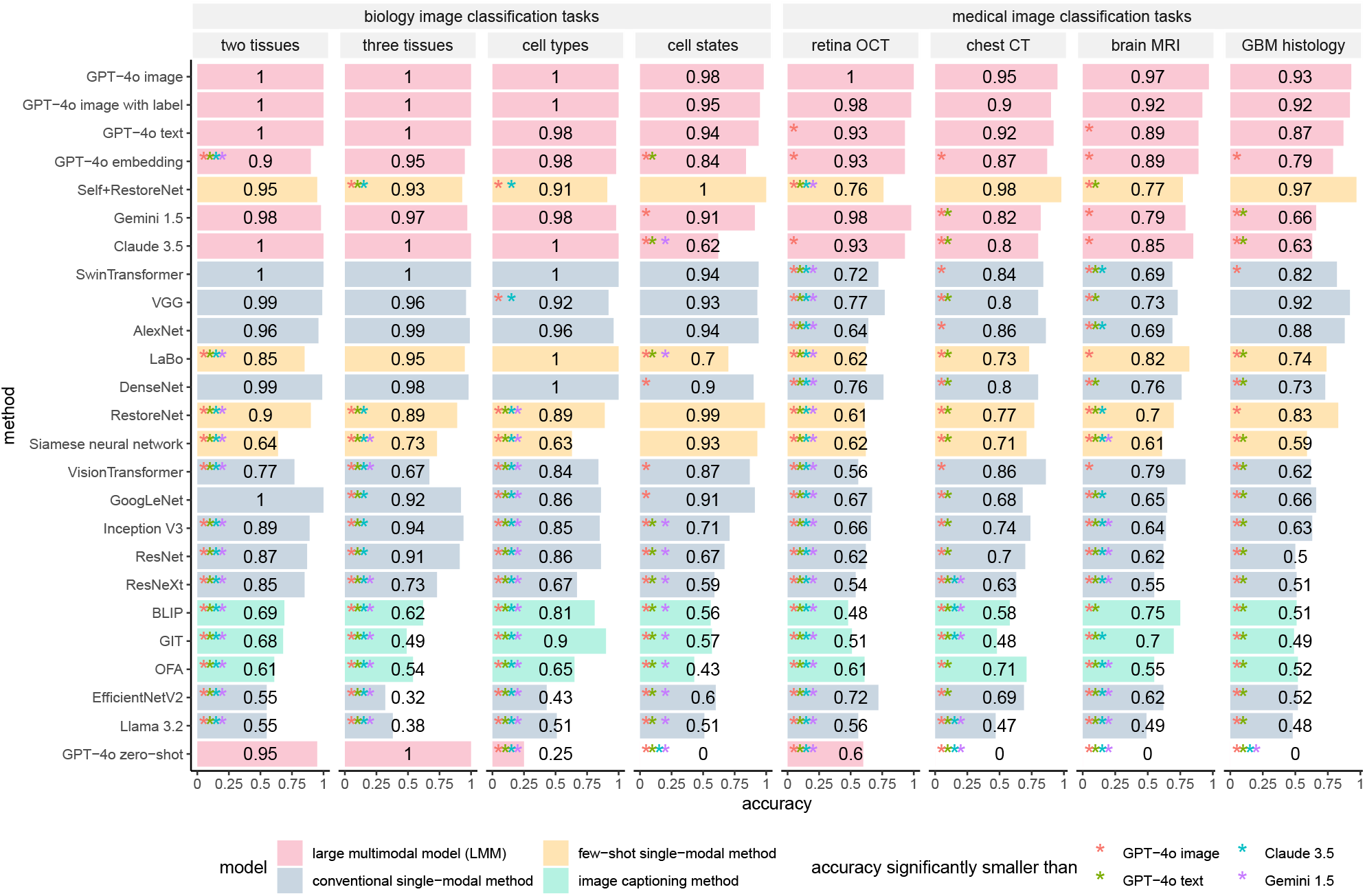
Accuracy of one-shot learning for LMMs and competing methods when training and testing images are homogeneous. “GPT-4o image”, “GPT-4o image with label”, “GPT-4o text”, “GPT-4o embedding”, and “GPT-4o zero-shot” refer to GPT-4o’s image-based classification, image-based classification where image category names are included in the training image array, text-based classification, embedding-based classification, and zero-shot classification, respectively. The performance of each method was compared to that of each LMM using a one-sided proportion test (R’s `prop.test()` function). The p-values were adjusted using the BH procedure^32^. Methods showing significant differences are marked with an asterisk (“*”), with colors corresponding to different LMMs.

We also tested two additional scenarios for GPT-4o. In the first scenario, the name of the image category is provided in the reference image. GPT-4o’s performance in this scenario (labeled as “GPT-4o image with label” in Figure 3) is highly consistent with the previous scenario described above (“GPT-4o image”), indicating that providing additional information about image category names does not improve the performance of image classification. In the second scenario, we evaluated GPT-4o’s zero-shot performance (labeled as “GPT-4o zero-shot” in Figure 3) by directly asking it to predict image categories without providing any training images (Methods). In this scenario, while GPT-4o performs almost perfectly in the two tissue classification tasks, it refuses to provide a definite answer in most other tasks, potentially due to legal concerns related to misclassification. Thus, GPT-4o’s zero-shot prediction may not be practical for real-world applications.

### LMMs outperform single-modal methods in generalizability

We further tested the models’ generalizability, which refers to their capacity to apply what they have learned from training images to systematically different testing images. While keeping the testing images unchanged, we introduced heterogeneity between the training and testing images by applying various transformations to the training images or by collecting training images from different studies (Methods, Figure 2).

Figure 4 shows the performance of all models when the training images and testing images are heterogeneous. Figure 5 further compares the performance of each model when the training images are either homogeneous or heterogeneous relative to the testing images. While the performance of all models decreases when the training and testing images are heterogeneous, the performance of LMMs declines much less than that of most single-modal models. Specifically, GPT-4o experiences only a slight performance drop, outperforming all single-modal methods in nearly all scenarios. The performance of SwinTransformer drops considerably in biology image tasks compared to its near-perfect performance when the training and testing images are homogeneous. A similar drop in performance can be observed for Self+RestoreNet. These results suggest that LMMs are more generalizable than single-modal methods and that LMMs can still reliably perform image classification tasks even when the training and testing images are heterogeneous.

**Figure 4.**
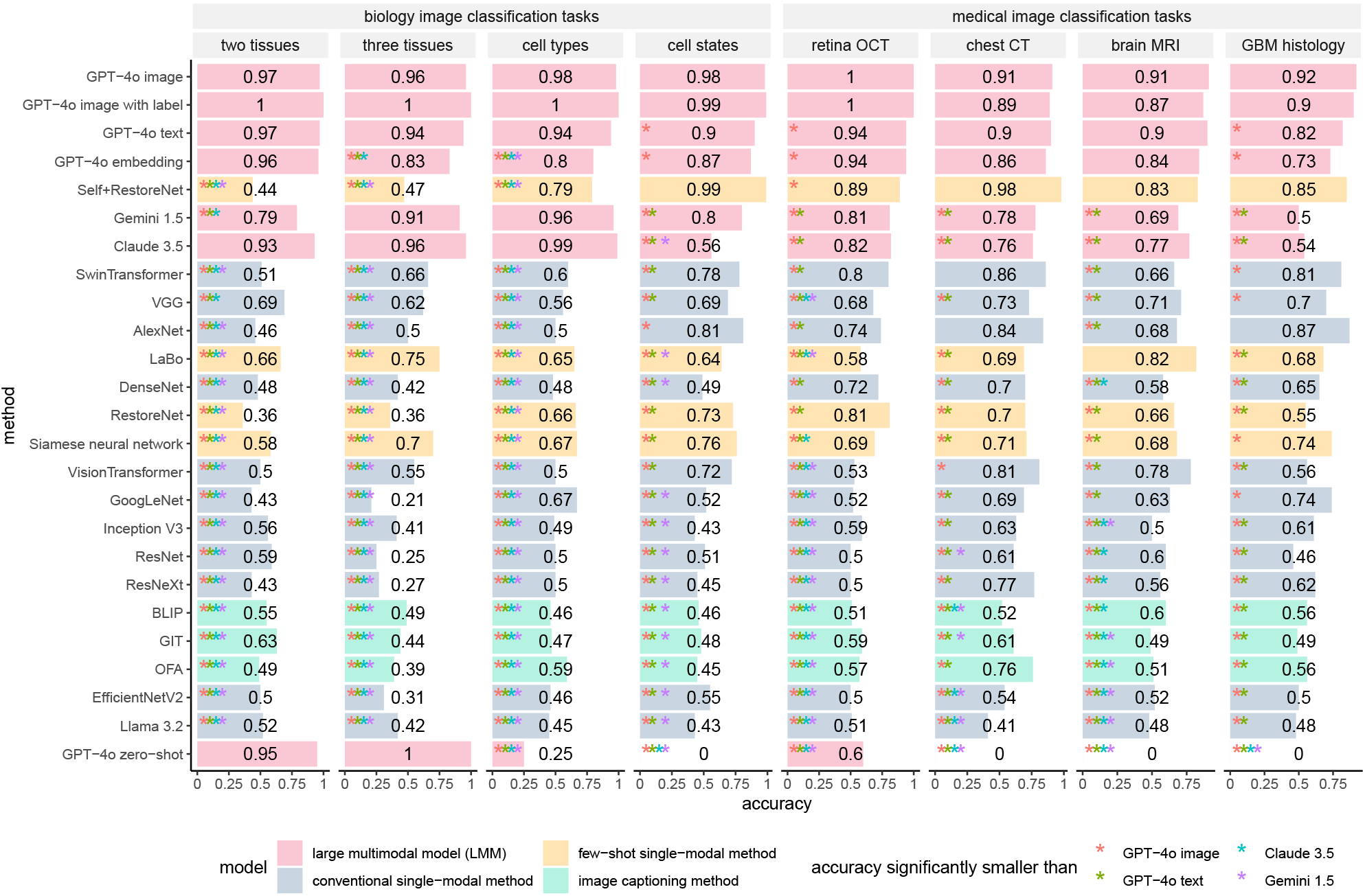
Accuracy of one-shot learning for LMMs and competing methods when training and testing images are heterogeneous. “GPT-4o image”, “GPT-4o image with label”, “GPT-4o text”, “GPT-4o embedding”, and “GPT-4o zero-shot” refer to GPT-4o’s image-based classification, image-based classification where image category names are included in the training image array, text-based classification, embedding-based classification, and zero-shot classification, respectively. The performance of each method was compared to that of each LMM using a one-sided proportion test (R’s `prop.test()` function). The p-values were adjusted using the BH procedure^32^. Methods showing significant differences are marked with an asterisk (“*”), with colors corresponding to different LMMs.

**Figure 5.**
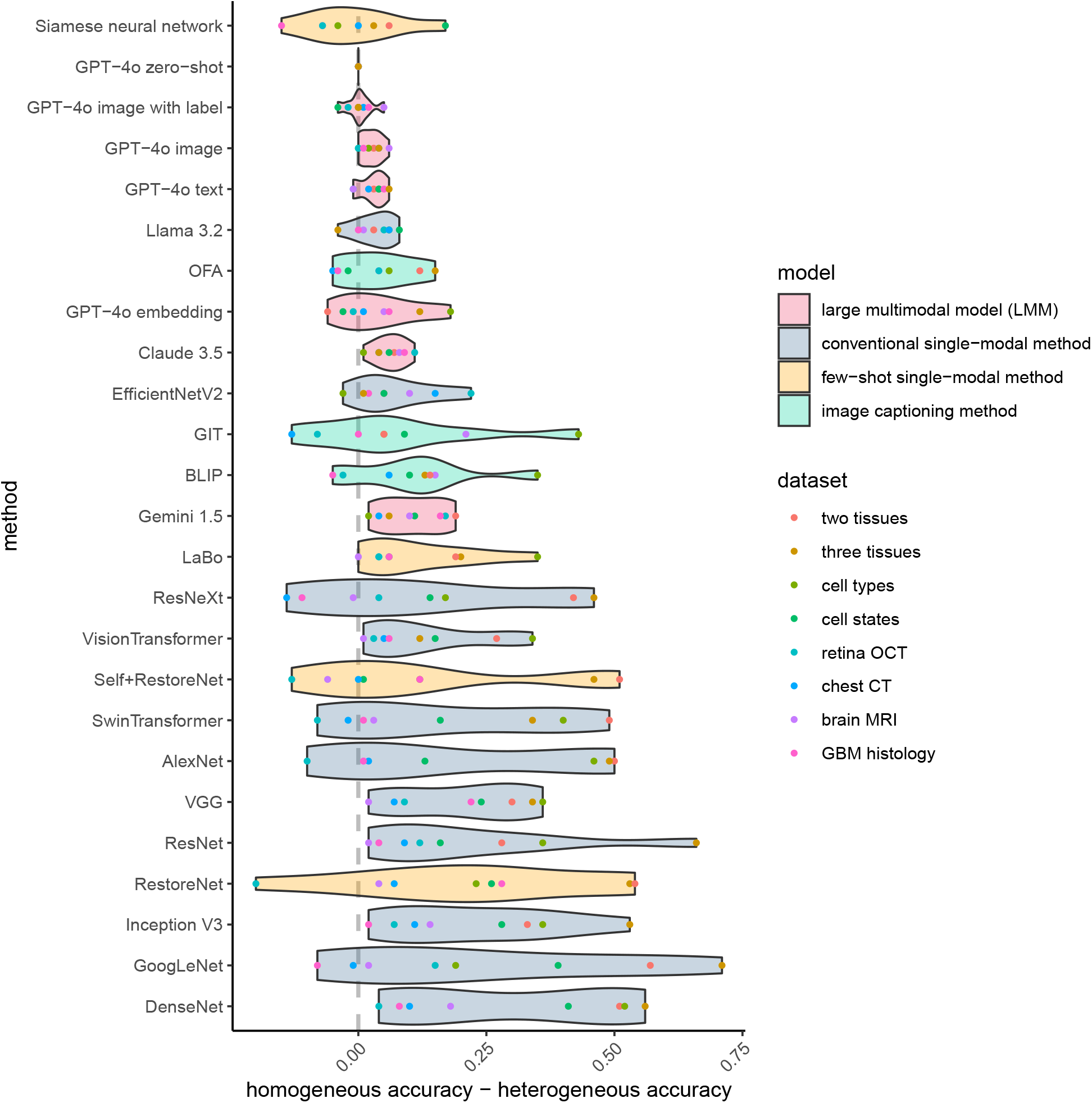
Differences in accuracy of each image classification method when training and testing images are homogeneous or heterogeneous.

### GPT-4o demonstrates interpretability

Beyond accuracy, we also evaluated the interpretability of GPT-4o, the best-performing LMM. We summarized the keywords from GPT-4o’s responses in image-based classification describing differences between the two image categories, and their enrichment is visualized as word clouds (Figure 6, Methods). The keywords align well with the features distinguishing the two image categories. For example, “dense” and “homogeneous” describe the image features of an artery, while “round” and “circular” describe those of a tibial nerve in tissue classification. In disease diagnostics, “uniform” and “smooth” describe the features of a normal retina, while “peak” and “irregular” describe those of DME. These results highlight GPT-4o’s interpretability in explaining the reasoning behind image classification, an ability not seen in single-modal methods. Additionally, the keywords provided by GPT-4o remain highly consistent regardless of the heterogeneity between the training and testing data, suggesting that GPT-4o can capture the most relevant information in image classification tasks, which may explain its superior generalizability. In contrast, single-modal methods can be misled by irrelevant information in the training data that does not transfer effectively to the testing data.

**Figure 6.**
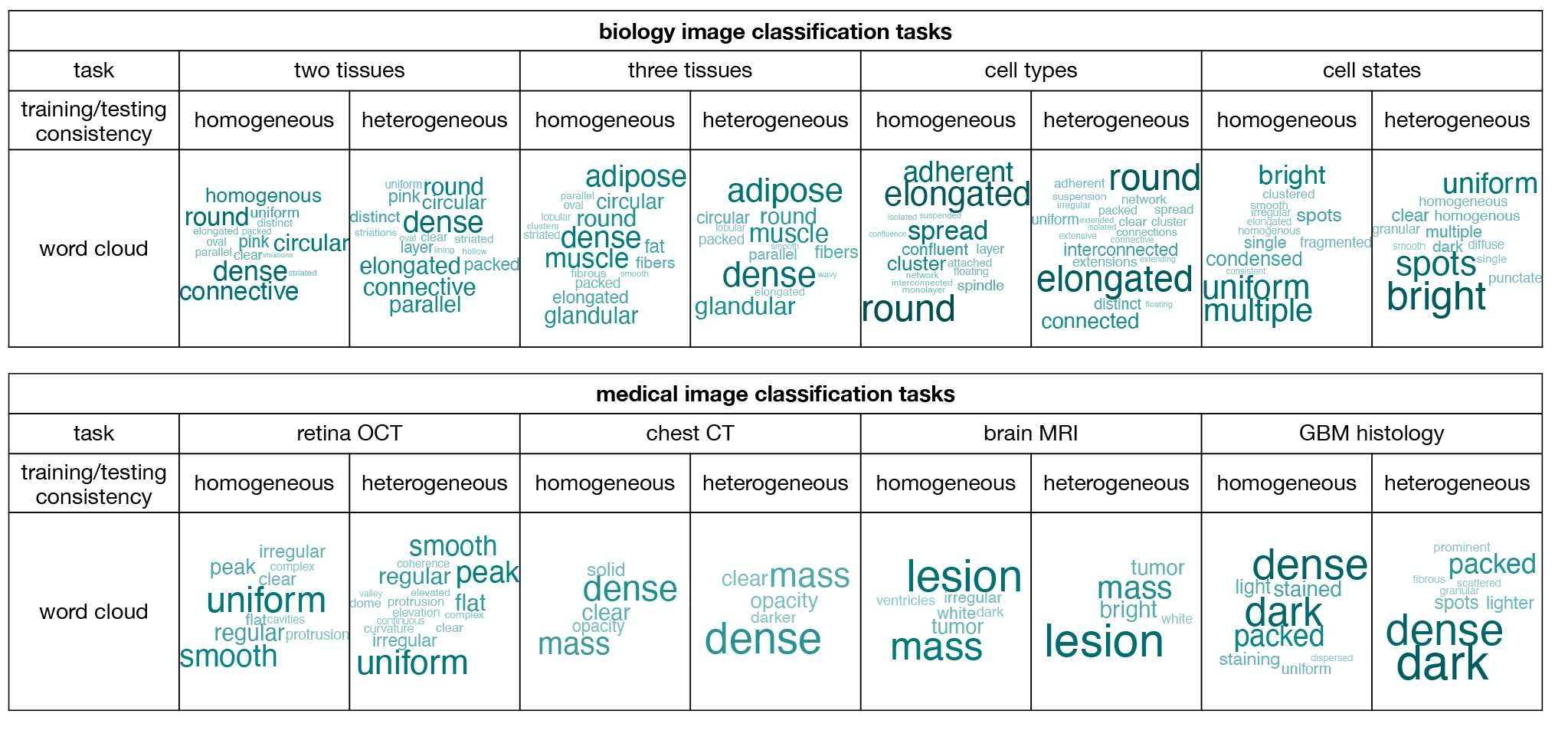
Word cloud showing keywords from GPT-4o’s responses in image-based classification.

We further studied the association between GPT-4o’s natural language descriptions in image-based classification and image features by computing image-text similarities (Methods). Using the CLIP model^10^, we mapped the natural language descriptions and images into embeddings within a shared space. For a pair of images from different categories, we calculated the cosine similarity of the differences between their image embeddings and text embeddings. In addition to these “matched” scenarios, we also designed “unmatched” scenarios, where the text embedding differences were generated from another pair of images belonging to the same category. Figure 7 presents the image-text similarity results for the two scenarios across each dataset. When text and images were matched, the image-text similarities were predominantly positive and significantly higher than those observed when text and images were mismatched in nearly all datasets. These findings further demonstrate GPT-4o’s ability to interpret image features as relevant natural language descriptions.

**Figure 7.**
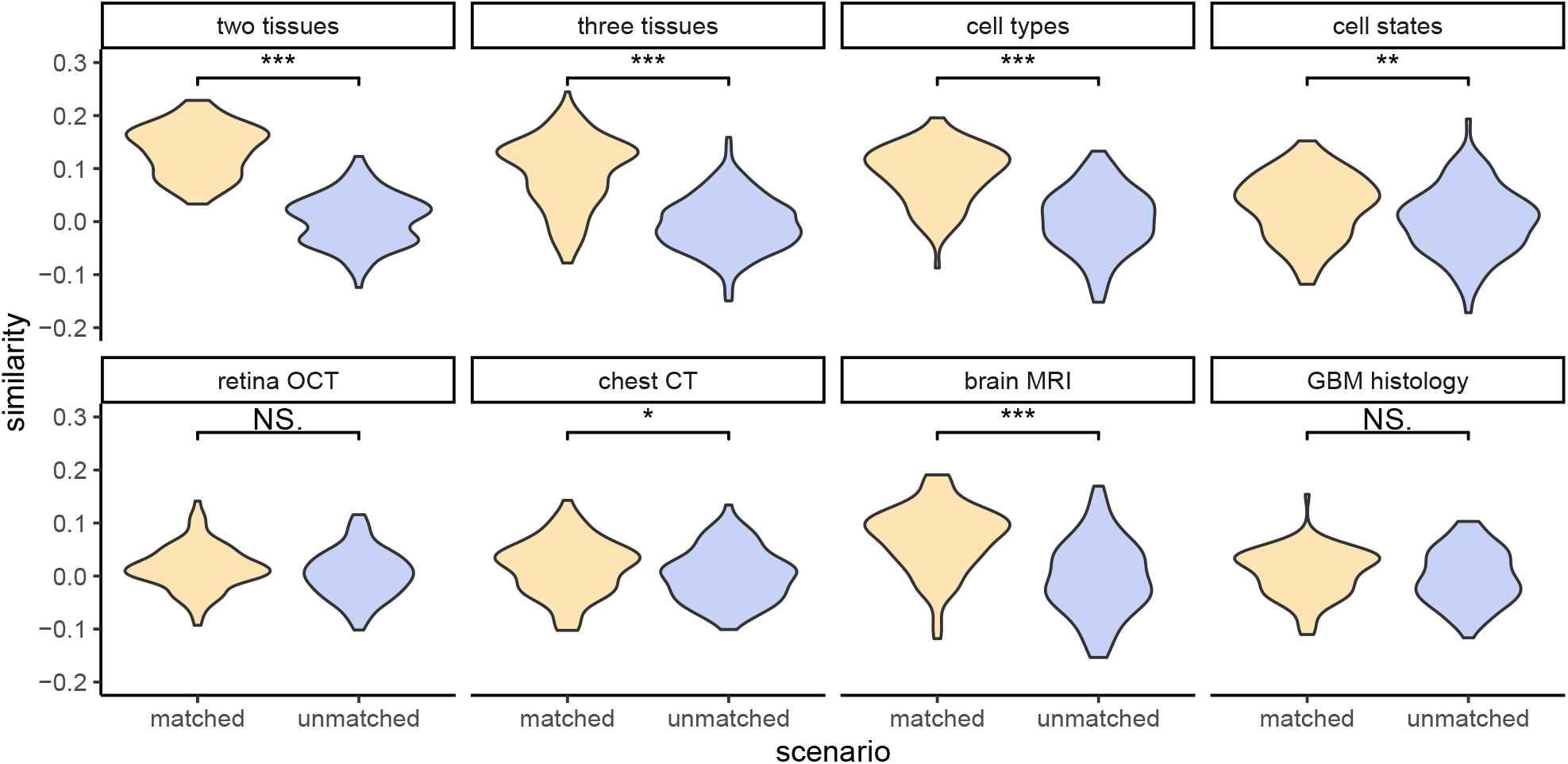
Image-text similarity for matched and unmatched pairs of images and natural language descriptions. A Wilcoxon test was performed to compare the similarity between matched and unmatched scenarios within each dataset. “***” indicates p-value < 0.001, “**” indicates p-value < 0.01 but > 0.001, “*” indicates p-value < 0.05 but > 0.01, and “NS.” indicates p-value > 0.05, denoting a statistically nonsignificant result.

### Text-based image classification with GPT-4o

GPT-4o’s interpretability inspired us to explore a new image classification paradigm using natural language cues. Such classification can be particularly useful when sharing training images is challenging due to technical or privacy concerns. We performed text-based and embedding-based image classification methods using GPT-4o, described in the previous section. The natural language that GPT-4o itself used to summarize the differences between image categories served as the sole information carrier for the training images. The performance of the two methods, labeled as “GPT-4o text” and “GPT-4o embedding” respectively in Figure 3, is only slightly worse than GPT-4o’s image-based classification (“GPT-4o image”) and outperforms all single-modal methods in almost all scenarios. The two classification methods also demonstrate superior generalizability, similar to its image-based classification (Figure 4,5). To better illustrate the embedding-based approach, we mapped the embeddings of images from different categories into a single UMAP space within each dataset (Figure 8). In most cases, the embeddings of images from the same category are clustered closely together, while those from different categories are well-separated. These results suggest that the text embeddings of natural language descriptions of image features effectively capture differences in image characteristics.

**Figure 8.**
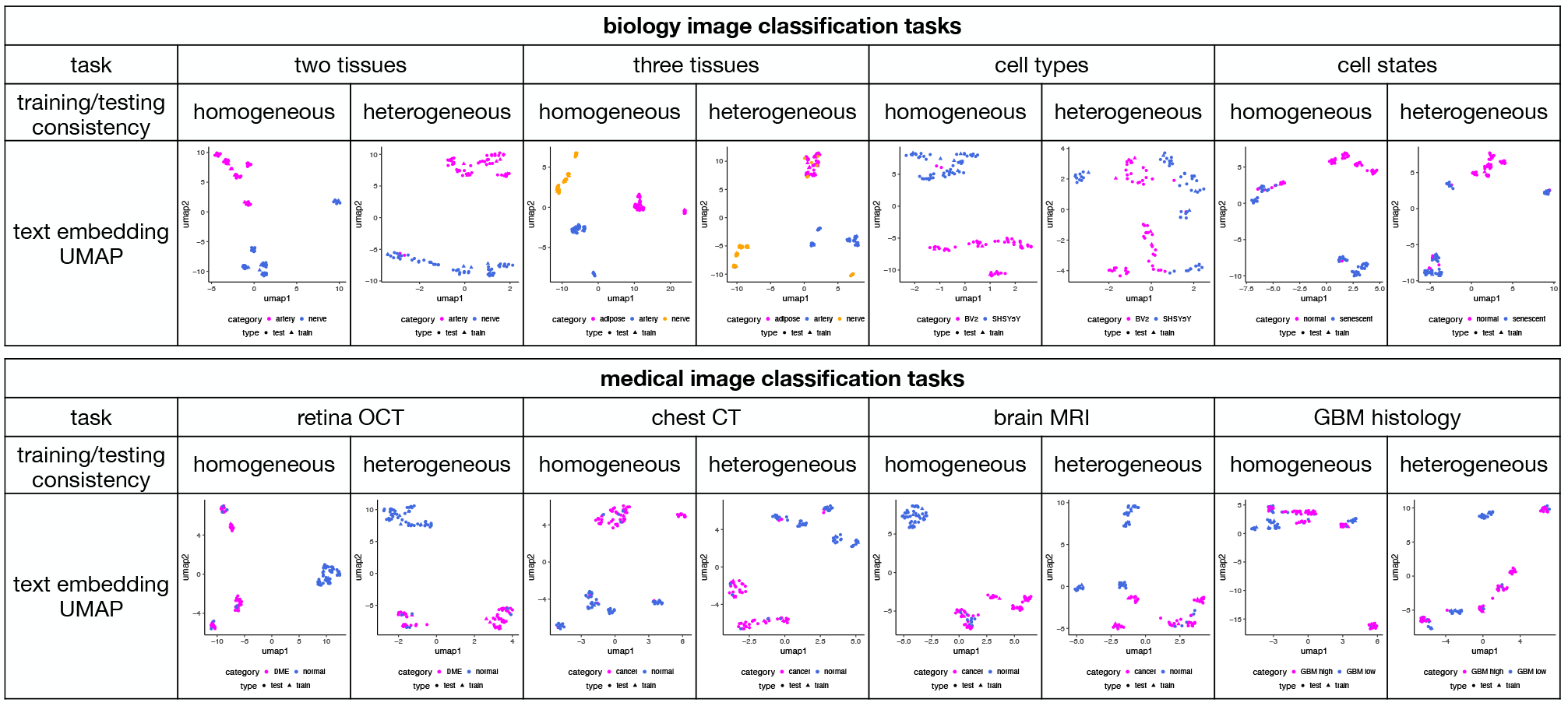
UMAP plots displaying the embeddings of natural language descriptions derived from image-based classification. Embeddings from different categories are distinguished by colors, while embeddings from training and testing images are represented with different shapes.

Note that embedding-based classification is also technically feasible using image captioning methods that do not rely on LMMs, such as BLIP^11^, GIT^12^, and OFA^13^. For comparison, we evaluated embedding-based classification using text captions generated by these approaches (Methods). The performance of these methods is substantially worse than GPT-4o’s embedding-based classification and also inferior to most single-modal methods. Consequently, GPT-4o is the only method that makes text-based image classification practical in real-world settings.

## Conclusions

Our study illustrates that LMMs, such as GPT-4o, are able to leverage natural language as a foundational component for reasoning in image classification. This feature substantially enhances LMMs’ capabilities in one-shot learning, generalizability, interpretability, and text-based image classification, surpassing single-modal methods. These findings indicate that LMMs, which integrate natural language with visual data, have potential advantages over single-modal methods that rely solely on images or image captioning methods that do not rely on LMMs. This insight could inform future research directions in computer vision, highlighting the benefits of combining linguistic and visual data. Moreover, the text-based classification approach suggests the possibility of developing new image analysis methods that utilize language-based communication between artificial intelligence systems. This approach could offer a more comprehensive way of interpreting and interacting with complex visual data, potentially enhancing the utility and applicability of AI in various domains by bridging the gap between human interpretative skills and machine efficiency.

Our study introduces several innovations to advance the understanding of LMMs in classifying biomedical images. First, we comprehensively evaluated the performance of LMMs and single-modal methods in one-shot image classification, clearly demonstrating the advantages of LMMs. We also compiled the images used in this study into a new benchmark dataset, bioimage1s, which can serve as a resource for future benchmarking studies. Second, we developed a framework for systematically evaluating the interpretability of LMMs. Using this framework, we assessed the interpretability of GPT-4o and compared its performance with conventional image captioning methods. Third, we designed two novel text-based image classification methods leveraging GPT-4o’s interpretability, thereby introducing a new paradigm for image classification. Our findings show that the performance of text-based classification is comparable to that of image-based classification.

It is important to note that the LMMs and single-modal methods evaluated in this study may rely on different sets of training data. For example, the conventional single-modal methods were trained on ImageNet-1K, a dataset with little overlap with the images evaluated in this study. In contrast, while the exact scope of the LMMs’ training data is proprietary and largely unknown, it is possible that LMMs were trained on a much more extensive dataset with greater overlap with the images evaluated here. To evaluate the performance of GPT-4o and single-modal methods under more comparable training data conditions, we conducted additional image classification tasks to differentiate between images of cats and dogs, as well as images of different types of dogs (Methods, Supplementary Figure 1). Both the training and testing images were directly obtained from the ImageNet-1K dataset, making them likely to have been included in the training data of GPT-4o and the conventional single-modal methods. The results demonstrate that GPT-4o remains the best-performing method. These findings suggest that the superior image classification performance of GPT-4o is likely not solely due to differences in training data but also attributable to the intrinsic performance of the model itself.

While LMMs offer powerful capabilities for biomedical image classification, their use comes with notable limitations. First, the financial cost of using commercial LMMs can be significant. For instance, platforms like GPT-4 and Claude require a monthly subscription fee (e.g., $20 for web access), and API usage costs scale with the volume of input and output tokens, making large-scale deployments expensive. Second, using these models raises data privacy concerns, as information provided in prompts may be collected by the companies operating these platforms. This poses a risk, particularly when sensitive biomedical data is involved. Third, the significant energy demands of training and operating LMMs contribute to environmental concerns, such as increased carbon emissions. Finally, there are ethical considerations, including potential biases in the training data, which can influence model predictions and perpetuate inequities. These challenges highlight the importance of balancing the benefits of LMMs with the need for ethical, equitable, and sustainable practices in their deployment.

## Methods

### Imaging data

#### Biology image classification tasks

For classification of two tissues, H&E images of artery and tibial nerve tissues were downloaded from the GTEx portal^14^. To create a scenario where training and testing images are homogeneous, we collected fifteen images of artery from different randomly chosen samples and fifteen images of tibial nerve from different randomly chosen samples. We randomly zoomed in on each of the original image provided by GTEx to display a part of the tissue, and all images have the same zoom level. We created five sets of training images, each set containing one artery image and one tibial nerve image. All the remaining images were used for testing. To create a scenario where the training and testing images are heterogeneous, we further zoomed in on each image of the same training set to display a smaller region of the tissue. The testing set remained the same.

For classification of three tissues, the same images of artery and tibial nerve tissues used for the classification of two tissues were utilized. Additionally, images of adipose tissue were collected from the GTEx portal and processed in the same manner.

For cell type classification, light microscopy images of BV-2 and SH-SY5Y cell lines were downloaded from the LIVECell dataset^15^. To create a scenario where training and testing images are homogeneous, fifteen images from each of the two cell lines were randomly chosen. Five images from each cell line were used to create five sets of training images, and the remaining images were used for testing. To create a scenario where the training and testing images are heterogeneous, we downloaded light microscopy images of BV-2 and SH-SY5Y from two additional studies^16,17^. For each image, we randomly cropped five different regions of the original image to obtain five new images for training. The testing set remained the same.

For cell state classification, DAPI fluorescence images were downloaded from a previous study^18^. To create a scenario where training and testing images are homogeneous, for each cell state of normal or senescence, fifteen images of single cells were randomly cropped from the original images in the study. Five images from each cell state were used to create five sets of training images, and the remaining images were used for testing. To create a scenario where the training and testing images are heterogeneous, we downloaded DAPI fluorescence images from another study^19^. For each cell state, five images of single cells were randomly cropped from the original images and five new sets of training images were created. The testing set remained the same.

#### Medical image classification tasks

For retina OCT classification, OCT images were downloaded from a previous study^20^. To create a scenario where training and testing images are homogeneous, fifteen images were randomly selected for each disease status. Five images for each disease status were used for training and the remaining images were used for testing. To create a scenario where the training and testing images are heterogeneous, we downloaded OCT images from another study^21^. For each disease status, five images were randomly selected for creating five sets of training images. The testing set remained the same.

For brain MRI classification, MRI images were downloaded from a Kaggle dataset (https://www.kaggle.com/datasets/navoneel/brain-mri-images-for-brain-tumor-detection). To create a scenario where training and testing images are homogeneous, fifteen images were randomly selected for each disease status. Five images for each disease status were used for training and the remaining images were used for testing. To create a scenario where the training and testing images are heterogeneous, we cropped either the left half or the right half of each training image. The testing set remained the same.

For chest CT classification, CT images were downloaded from a Kaggle dataset (https://www.kaggle.com/datasets/mohamedhanyyy/chest-ctscan-images). To create a scenario where training and testing images are homogeneous, fifteen images were randomly selected for each disease status. Five images for each disease status were used for training and the remaining images were used for testing. To create a scenario where the training and testing images are heterogeneous, we applied a horizontal flip to each training image. The testing set remained the same.

For GBM histology classification, histology H&E images from glioblastoma (GBM) patients were downloaded from The Cancer Imaging Archive^22^ via the Clinical Proteomic Tumor Analysis Consortium (CPTAC) portal. To create a scenario where the training and testing images are homogeneous, fifteen images were randomly selected from samples with either at least 90% tumor nuclei (high GBM density) or at most 25% tumor nuclei (low GBM density). Five images from each GBM density group were used for training, and the remaining images were used for testing. To simulate a scenario where the training and testing images are heterogeneous, the original color training images were converted to black-and-white images, while the testing set remained unchanged.

#### Animal classification tasks

For the classification of cats and dogs, images of cats (ID: n02123394), the Komondor dog (ID: n02105505), and the Alaskan Malamute dog (ID: n02110063) were downloaded from the ImageNet 1K dataset. For the cat images, five were randomly selected for training, and ten were randomly selected for testing. For the Komondor dog images, two were randomly selected for training, and five were randomly selected for testing. For the Alaskan Malamute dog images, three were randomly selected for training, and five were randomly selected for testing. Images of the Komondor dog and the Alaskan Malamute dog were both treated as images of dogs.

For the classification of dog types, the same images of the Komondor dog and the Alaskan Malamute dog from the ImageNet 1K dataset were used. For each dog type, five images were randomly selected for training, and ten images were randomly selected for testing.

### Image classification with LMMs

#### Specifics of LMMs

Image classification tasks with LMMs were all performed using the application programming interfaces (APIs) provided by the companies that developed them.

GPT-4o was performed with the `gpt-4o-2024-08-06` model via the `openai` python package (version 1.42.0) provided by OpenAI.

Claude 3.5 Sonnet was performed with the `claude-3-5-sonnet-20240620` model via the `anthropic` python package (version 0.34.2) provided by Anthropic.

Gemini 1.5 was performed with the `gemini-1.5-flash` model via the `google.generativeai` python package (version 0.8.0) provided by Google.

Llama 3.2 11B vision model was performed with the `Llama-3.2-11B-Vision-Instruct` model via the `transformers` python package (version 4.47.1) provided by Hugging face.

#### Image-based classification

Image-based classification was performed for all LLMs. An LMM was given a reference image array consisting of two training images as the first image, the testing image as the second image, and queried with the following prompt message for tissue classification:

“There are 2 rows in total for the first image. Each row represents example images from a tissue. The second image is another image of a tissue. Which row in the first image is more similar to the second image and why?”

For cell type classification, cell state classification, retina OCT, chest CT, and brain MRI, the word “tissue” was replaced with “cell type”, “cell state”, “status of a tissue”, “lung abnormality”, and “brain abnormality” respectively in the previous prompt message.

For GPT-4o, two additional strategies were implemented. In the first strategy (referred to as “GPT-4o image with label” in Figure 3), the labels in the first column of the training image array were replaced with the names of the image categories. Image classification was then performed using the same prompts as above. In the second strategy (referred to as “GPT-4o zero-shot” in Figure 3), GPT-4o was only provided with the test image, and the following prompt was used: “Is this image cancer or normal?” Here, “cancer” and “normal” were replaced with the corresponding names of the image categories for each image classification task.

#### Text-based classification

Text-based classification was only performed for GPT-4o. GPT-4o was first given a reference image array consisting of two training images and queried with the following prompt message:

“What are the most distinct features differentiating the two rows of images? Provide detailed explanations.”

The response by GPT-4o was recorded, and attached to the bottom of the following sentence to form a new prompt message. “Is the given image more consistent with row 1 or row 2 based on the descriptions below?”

In a completely new conversation, GPT-4o was then given the testing image and queried with the above new prompt message.

#### Embedding-based classification

Embedding-based classification was performed only for GPT-4o. We first manually curated GPT-4o’s description of each training and testing image from its natural language responses during text-based classification. Each image description was converted into a numeric vector of text embeddings using the “text-embedding-3-small” model provided by OpenAI. For every pair of training images and each testing image, a cosine similarity was calculated between the testing image and each training image. The category of the training image with the highest cosine similarity was assigned as the predicted category of the testing image.

To visualize the embeddings in UMAP space, the embedding vectors of training and testing images from different image categories were combined into a single matrix. The UMAP space was generated using the `umap` function with default settings from the `UMAP` R package (version 0.2.10.0).

### Competing methods

#### Conventional single-modal methods for image classification

The Python `torchvision` package^23^ (version 0.14.1), part of the `PyTorch` project^24^, was used to implement the ten single-modal methods for image classification with pre-trained weights. Specifically, the following models and their pre-trained weights were used: `alexnet` model with `AlexNet_Weights`, `densenet201` model with `DenseNet201_Weights`, `efficientnet_v2_m` model with `EfficientNet_V2_M_Weights`, `googlenet` model with `GoogLeNet_Weights`, `inception_v3` model with `Inception_V3_Weights`, `resnet50` model with `ResNet50_Weights`, `resnext101_32×8d` model with `ResNeXt101_32×8D_Weights`, `swin_b` model with `Swin_B_Weights`, `vgg16` model with `VGG16_Weights`, and `vit_b_32` model with `ViT_B_32_Weights`. The pretrained models were fine-tuned with training images provided in this study with a cross entropy loss, Adam optimizer^25^, and a learning rate of 0.0001. The model training was stopped when the training loss is smaller than 0.00001 or when the training exceeds 50 epochs, whichever happens first.

#### Few-shot single-modal methods for image classification

##### Siamese Neural Network

To implement the Siamese neural network, we adopted the architecture recommended in the original paper^26^ and trained the model on the miniImageNet dataset, a widely recognized benchmark for one-shot learning tasks^27^. We applied the same loss function and set the learning rate to 0.0001. The pre-trained model was then used as the initialization for fine-tuning on the training images provided in our study. For each classification task, we augmented the training images by applying horizontal flipping and rotations of 90°, 180°, and 270°. To construct image pairs for testing, we iterated over all positive and negative pairs in our test sets. Positive pairs consisted of images from the same category, while negative pairs consisted of images from different categories. Fine-tuning was performed on the augmented training images with a learning rate ranging from 0.001 to 0.0001, adjusted based on performance. The fine-tuning process was terminated when the training loss dropped below 0.05 or after 50 epochs, whichever occurred first.

##### RestoreNet and Self+RestoreNet

The RestoreNet and Self-RestoreNet models^27^ trained on the miniImageNet dataset were utilized. Image datasets in this study were treated as the novel set, with the training images serving as the support set and the testing images as the query set. We evaluated both versions of the proposed models described in the original paper: one with prototype transformation only (RestoreNet) and another combining prototype transformation with self-training (Self-RestoreNet). The skip-connection rate *p* was varied across [0.05, 0.95] to evaluate the model performance.

##### LaBo

To apply LaBo^28^, we designed prompts tailored to each classification task during the concept generation step. For example, in the two-tissue classification task, we used prompts such as:: “Describe what the tissue type of artery/tibial nerve in H&E images looks like.”, “Describe the appearance of the tissue type of artery/tibial nerve in H&E images.”, “Describe the pattern of the tissue type of artery/tibial nerve in H&E images.”, “Describe the shape of the tissue type of artery/tibial nerve in H&E images.” For concept selection, we used submodular optimization^29^ to select 10 concepts for each class. For concept-image alignment, we used the ViT-L/14 model to obtain concept and image embeddings. The weight matrix *W* was initialized using language model priors from GPT-4. Specifically, if a concept *c* was associated with a class *y*, the corresponding element *W*_*y,c*_ was initialized to 1 before optimization; otherwise, it was set to 0. Training of *W* was conducted using cross-entropy loss, the Adam optimizer^25^, and a learning rate in the range [10^−1^, 10^−5^]. The training process stopped when the loss dropped below 0.00001 or after 50 epochs, whichever occurred first.

#### Image captioning methods

##### BLIP

The pretrained Salesforce/blip-image-captioning-base model was utilized using the `transformers` library provided by Hugging Face. Captions were generated by processing images and textual prompts, with a maximum length of 20 tokens, using greedy decoding (num_beams=1). A repetition penalty of 1.0 was applied to prevent redundant output, and early stopping ensured that caption generation halted at the first complete result.

##### OFA

We utilized pretrained models through both Fairseq and Hugging Face implementations. The Fairseq version employed patch-based image encoding with manual generation, while the Hugging Face version leveraged integrated generation functions. Beam search (num_beams=5) and n-gram constraints (no_repeat_ngram_size=3) were applied to the Hugging Face model to enhance caption diversity. The same tokenizer was used in both approaches to ensure consistency during evaluation.

##### GIT

We utilized the pretrained microsoft/git-large model via the Hugging Face `transformers` library. Captions were generated by encoding images with the AutoProcessor and running the model with a maximum length of 50 tokens. Beam search (num_beams=5) and n-gram constraints (no_repeat_ngram_size=3) were applied to minimize repetition. This configuration ensured consistent and high-quality caption generation across all images.

### Performance evaluations for image classification

Each classification result from the LMMs was manually evaluated for accuracy by a human annotator and verified by another. A score of 0 was given if the answer was inaccurate or if the LMM refused to provide a definite answer. The classification results from other methods were evaluated programmatically.

For an image classification task, an image classification method was trained on each of the five reference image arrays and tested on each of the 20 testing images, resulting in 100 predicted results. The accuracy is defined as the percentage of the 100 predicted results that agree with the correct answers.

More formally, denote *p*_*i j*_ as the predicted binary label when a method was trained on *i*th (*i* = 1, 2, 3, 4, 5) reference image array and tested on *j*th (*j* = 1, …, 20) testing image. Denote *l* _*j*_ as the true binary label for *j*th testing image. The accuracy is defined as ∑_*i, j*_(*p*_*i j*_ = *l* _*j*_)*/*100. The R package `ggplot2` (v.3.3.0)^30^ was used for visualizing the performance.

### GPT-4o word cloud

For each classification task, we calculated the number of times each word occur in the 100 responses by GPT-4o, after removing special characters. A word cloud is created using the `wordcloud` R package (version 2.6)^31^ for words that occur in at least 10% of the 100 GPT-4o responses. Words without specific meanings (e.g. articles such as “a” and “the” and conjunctions such as “and”) were removed manually.

### Image-text similarity analysis

For each testing image and its corresponding natural language descriptions generated by GPT-4o, we convert them into embeddings in a shared space using the CLIP model^10^ (clip-vit-base-patch32) via the `transformers` package. Note that each testing image is evaluated five times, as there are five sets of training images. Consequently, five sets of natural language descriptions are generated by GPT-4o for each testing image.

In a scenario where the images and text descriptions are matched, we randomly selected a pair of images from different image categories. We then obtained their natural language descriptions from one randomly chosen round of the five evaluations. A cosine similarity was calculated between the differences in the embeddings of the two images and the differences in the embeddings of their natural language descriptions. This entire process was repeated 100 times.

In a scenario where the images and text descriptions are unmatched, the procedure is the same as in the matched scenario, except that the natural language descriptions were obtained from a pair of randomly selected images belonging to the same image category.

## Acknowledgments

Z.J., H.M., and Y.Q. were supported by the National Institutes of Health under Award Number U54AG075936 and R35GM154865. W.H. and Q.L. were supported by the National Institute of General Medical Sciences of the National Institutes of Health under Award Number R35GM150887 and the General Fund at Columbia University Department of Biostatistics. The content is solely the responsibility of the authors and does not necessarily represent the official views of the National Institutes of Health.

## Author contributions

W.H. and Z.J. conceived the study. W.H., Q.L., H.M., Y.Q., and Z.J. conducted the analysis. W.H. and Z.J. wrote the manuscript.

## Data availability

H&E images for tissue classification were downloaded from the GTEx Histology Viewer of the GTEx portal^14^ (https://gtexportal.org/home/histologyPage). Light microscopy images for cell type classification were downloaded from previous studies^15–17^. DAPI fluorescence images for cell state classification were downloaded from previous studies^18,19^. Retina OCT images for disease diagnostics were downloaded from previous studies^20,21^. Brain MRI images were downloaded from a Kaggle dataset (https://www.kaggle.com/datasets/navoneel/brain-mri-images-for-brain-tumor-detection Chest CT images were downloaded from a Kaggle dataset (https://www.kaggle.com/datasets/mohamedhanyyy/chest-ctscan-images). GBM histology images were downloaded from The Cancer Imaging Archive CPTAC Pathology Portal (https://www.cancerimagingarchive.net/collection/cptac-gbm/). Responses of LMMs in this study are available as supplementary materials. All training and testing images are available at GitHub (https://github.com/zji90/bioimage1s).

## Code availability

All codes to reproduce the presented analyses are publicly available in Github repository https://github.com/Winnie09/gptimage.

