## Supplementary Figure 1 for "Assessing large multimodal models for one-shot learning and interpretability in biomedical image classification"

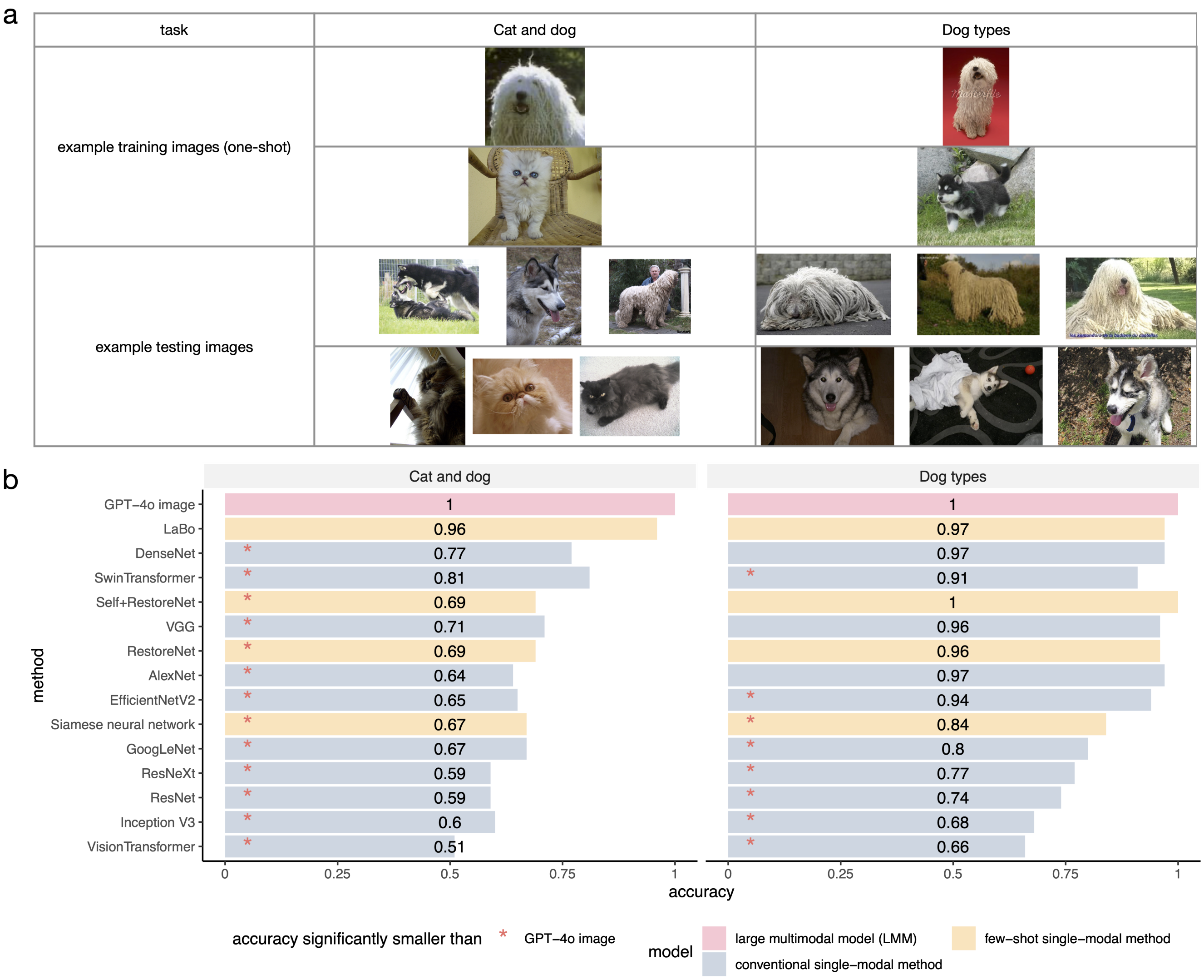


**Supplementary Figure 1**: Classification of images of cats and dogs, as well as images of different dog types. **a**, Example training and testing images for each classification task. **b**, Accuracy of one-shot learning for LMMs and competing methods. "GPT-4o image" refers to GPT-4o's image-based classification. The performance of each method was compared to GPT-4o using a one-sided proportion test (R's `prop.test()` function). The p-values were adjusted using the BH procedure. Methods showing significant differences are marked with an asterisk ("*").
